## Supplemental Data (Figures) for "Epigenetics and chromatin structure regulate *var2csa* expression and the placental binding phenotype in *Plasmodium falciparum*"

^†^ Joint authors

**This document includes the following:**

Figures S1 to S8

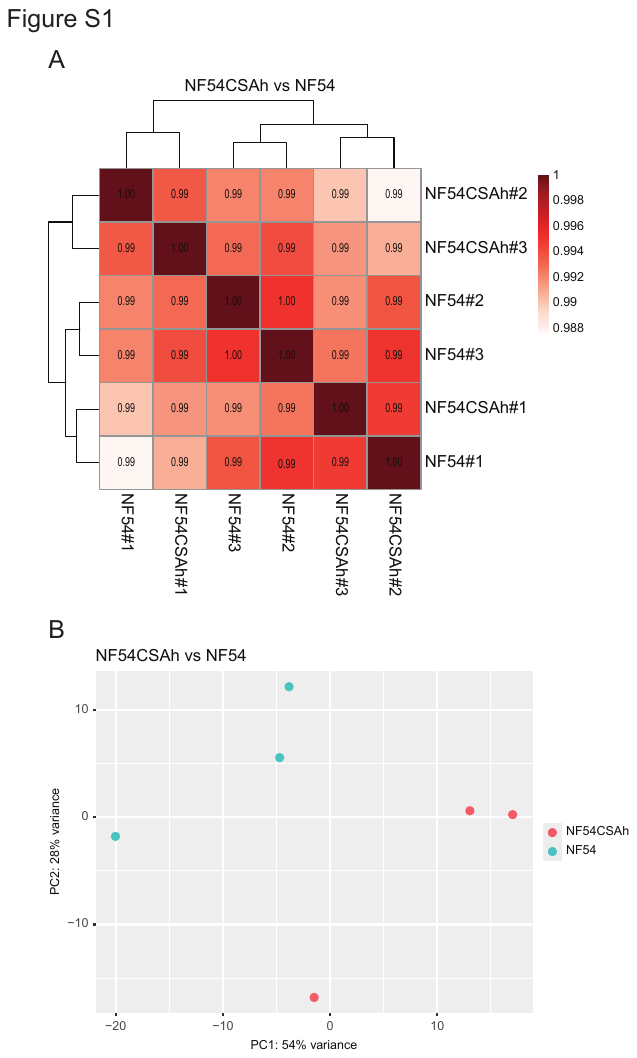

**Figure S1.** Genome-wide expression profiles are highly similar among all samples. (**A**) Correlation heatmap of regularized logarithmic read counts displaying strong similarity in expression profiles between replicates and samples. (**B**) Principal component analysis shows similar variance between replicates of each sample.

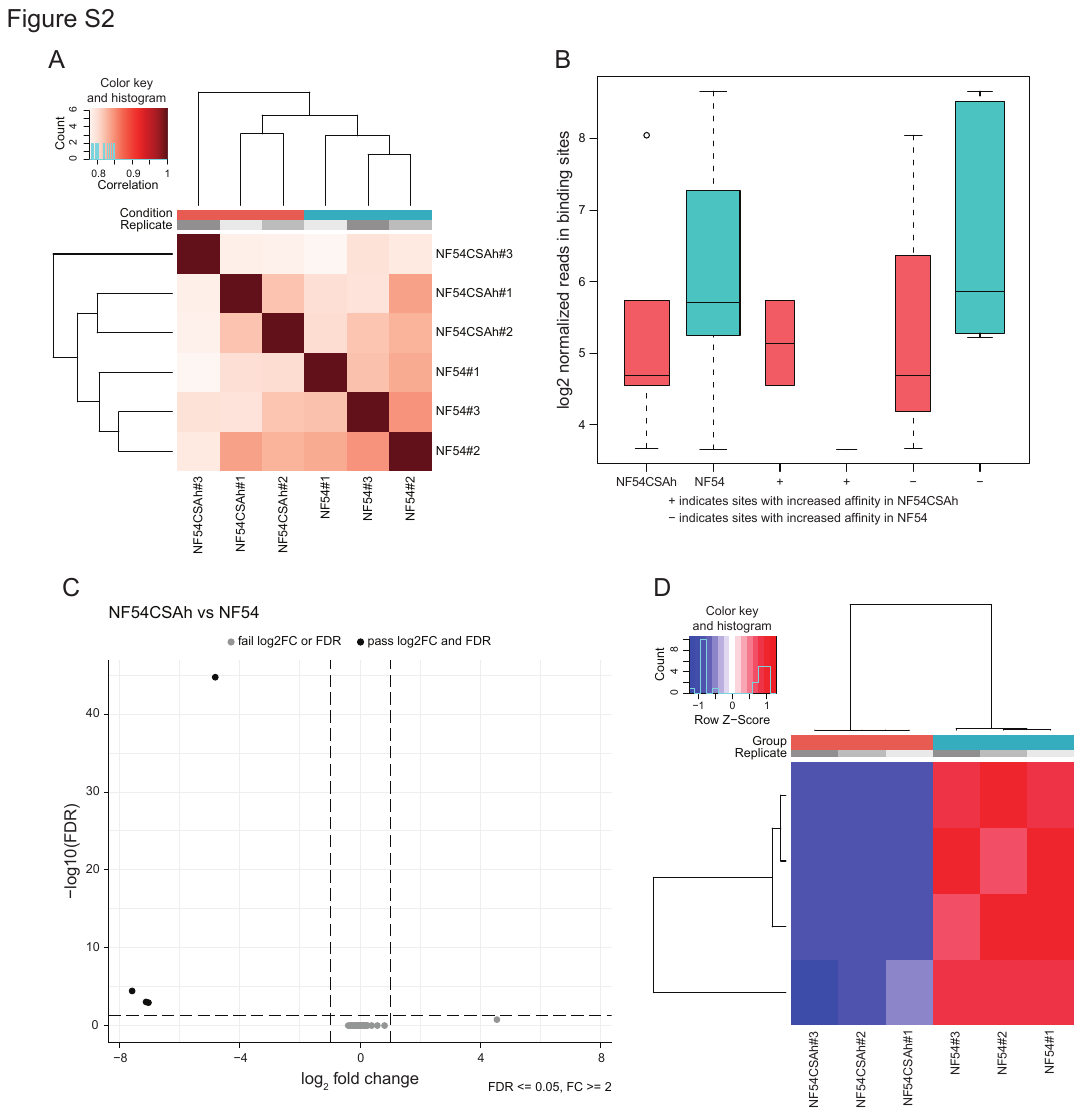

**Figure S2.** Differential peak calling of H3K9me3 reveals highly focused changes in *var2csa* region. (**A**) Correlation analysis indicates similarity in global binding of H3K9me3 between replicates and samples. (**B**) Read counts are greater in NF54 than NF54CSAh within all peaks identified by differential binding (FDR < 0.05) analysis. (**C**) There are only four significantly differentially bound (FDR < 0.05) sites between NF54 and NF54CSAh with a log₂ fold change greater than 1 (black)**,** all of which contain more reads in NF54 than NF54CSAh. (**D**) Read counts for the four differentially bound regions are consistent among replicates within each sample.

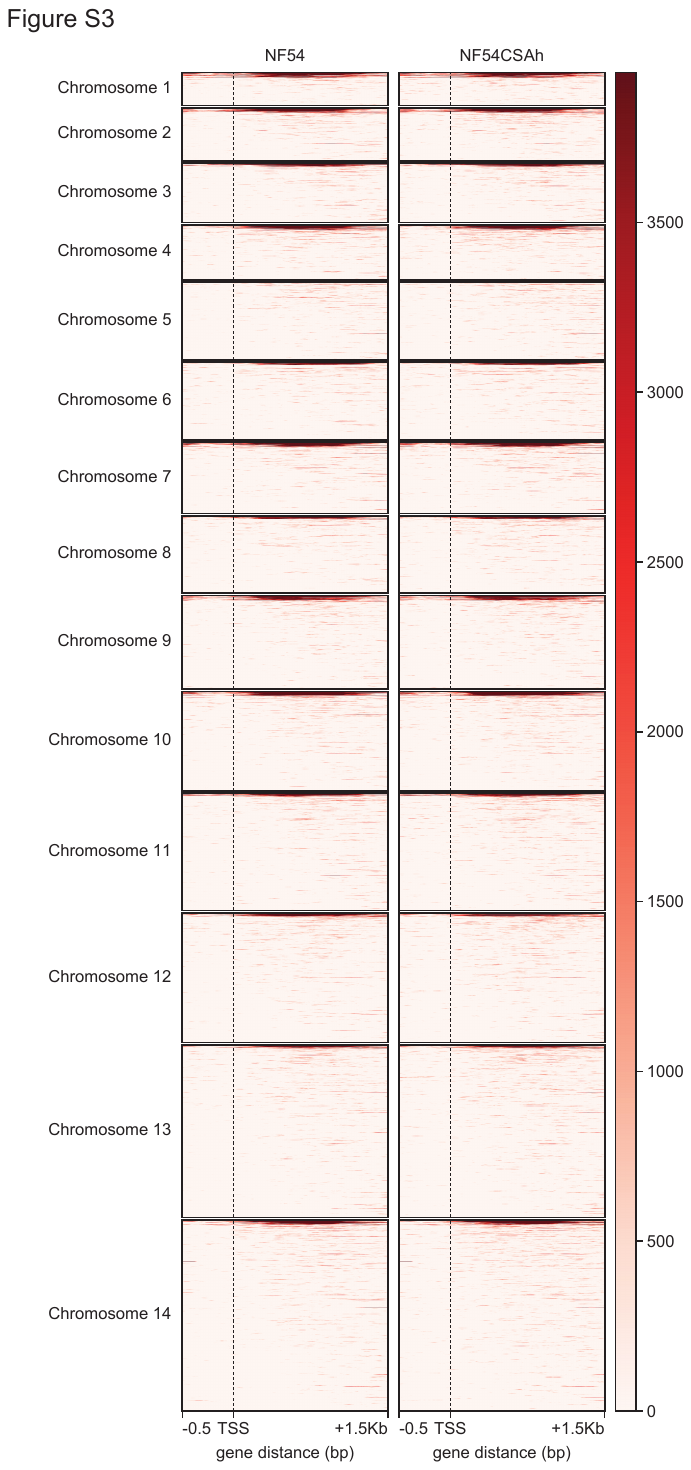

**Figure S3.** A majority of genes across all chromosomes contain low levels of H3K9me3 surrounding the TSS. Heatmaps of input-normalized H3K9me3 occupancy for all genes separated by chromosome in NF54 and NF54CSAh (0.5 kb 5’ of the transcription start site (TSS) and 1.5kb 3’ of the TSS). Genes within each chromosome are sorted by the mean read count within the 2 kb region surrounding the TSS.

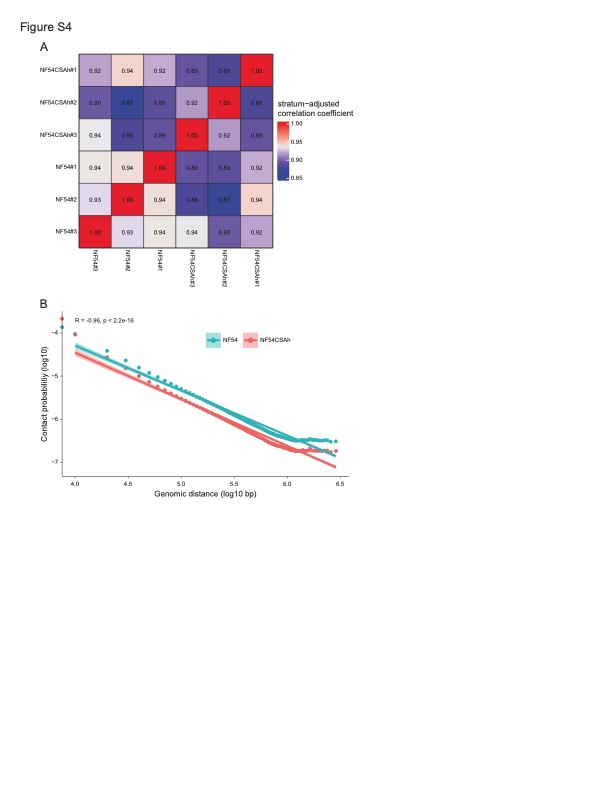

**Figure S4.** There is a strong correlation in genome-wide interactions between samples and replicates and between contact count probability and genomic distance. (**A**) Distance normalized correlation heatmap showing similar correlation between replicates of both samples. (**B**) Scatter plot indicating a negative log-linear relationship between contact probability and genomic distance for NF54 and NF54CSAh.

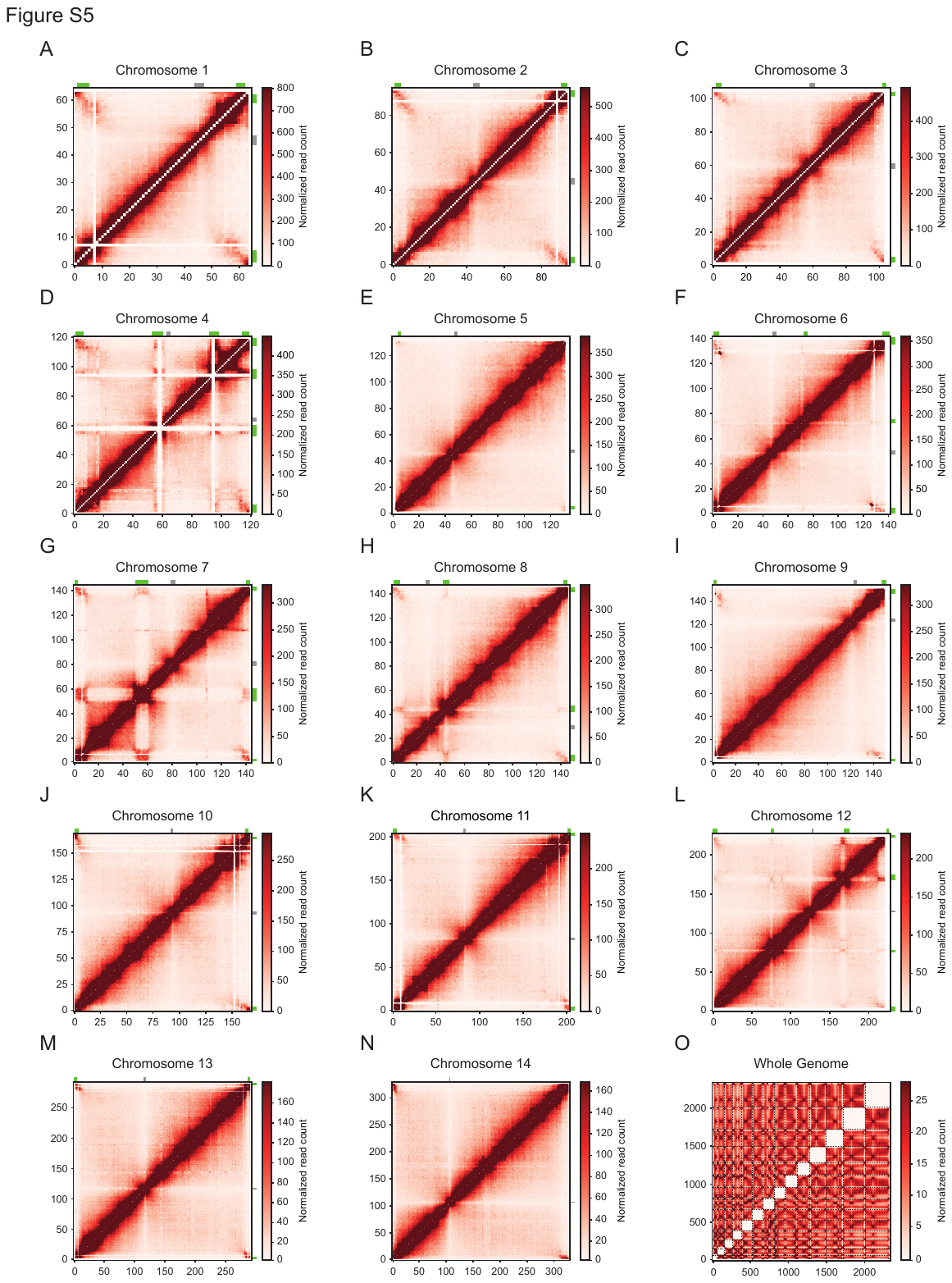

**Figure S5.** Hi-C intrachromosomal and interchromosomal contact count heatmaps for NF54. (**A-M**) Heatmaps generated from ICED (26619908) and per-million read count normalized intrachromosomal contact count matrices binned at 10-kb resolution for all 14 chromosomes of NF54 with annotated centromeres (gray) and *var* gene containing bins (green). (**N**) Heatmap depicting the normalized genome-wide interchromosomal contact count matrix for NF54 with intrachromosomal interaction matrices removed.

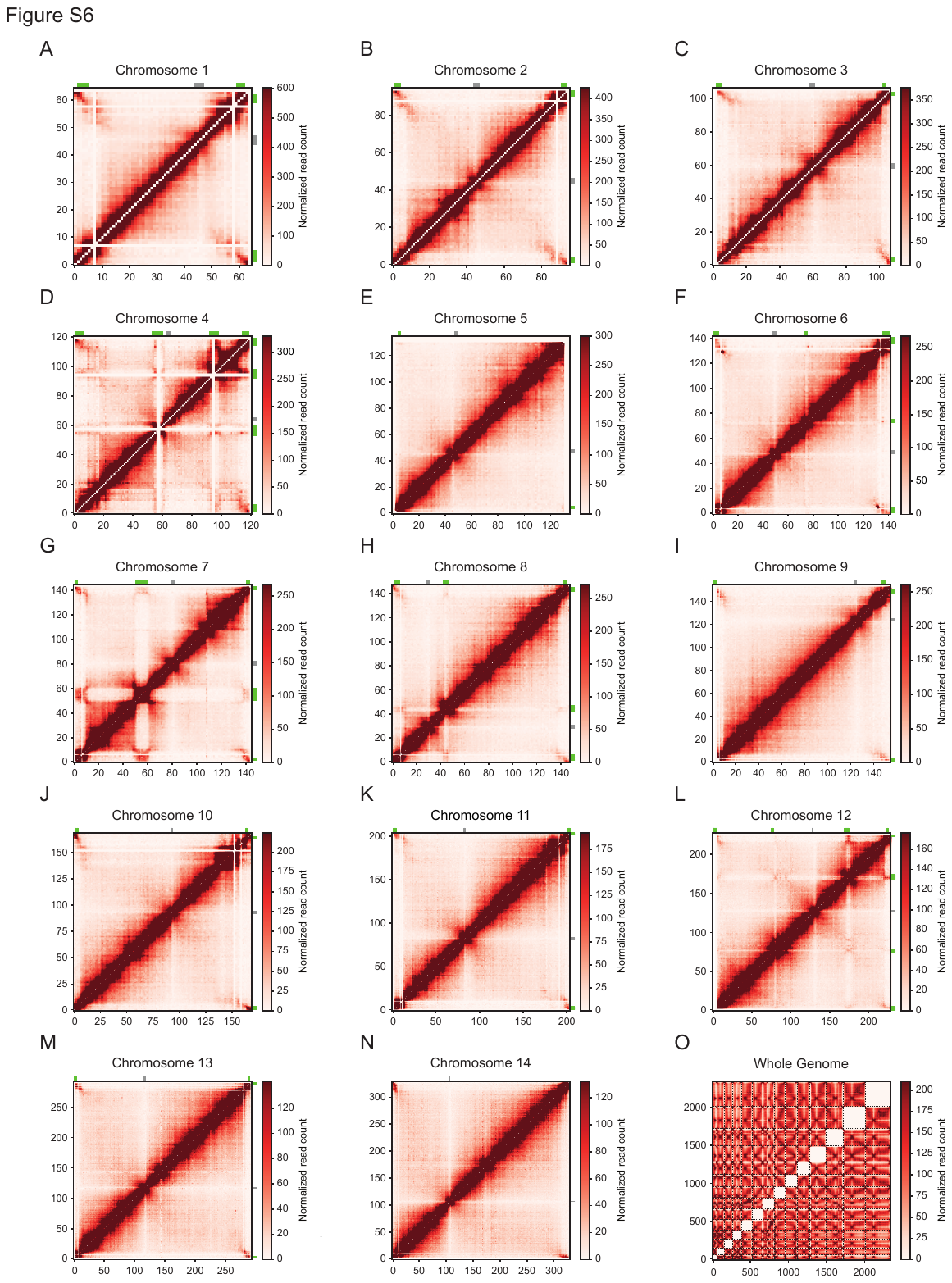

**Figure S6.** Hi-C intrachromosomal and interchromosomal contact count heatmaps for NF54CSAh. (**A-M**) Heatmaps generated from ICED (26619908) and per-million read count normalized intrachromosomal contact count matrices binned at 10-kb resolution for all 14 chromosomes of NF54CSAh with annotated centromeres (gray) and *var* gene containing bins (green). (**N**) Heatmap depicting the normalized genome-wide inter-chromosomal contact count matrix for NF54CSAh with intrachromosomal interactions removed.

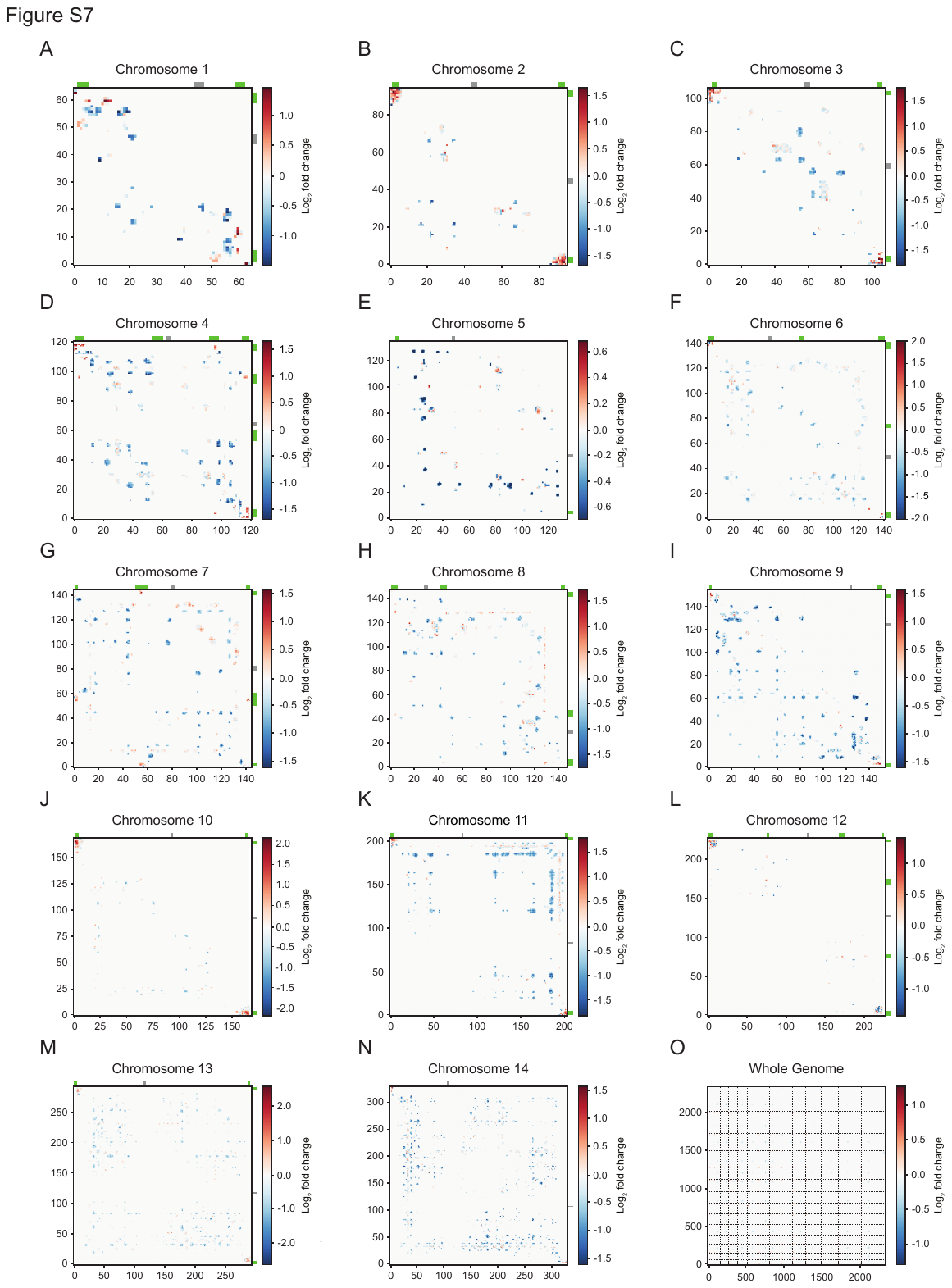

**Figure S7.** Differential interaction contact count heatmaps show higher interactions between telomeric and internal *var* gene-containing regions in NF54CSAh. (**A-M**) Heatmaps generated from differential interaction matrices for all 14 chromosomes, identifying regions with an increase (red) or decrease (blue) of intrachromosomal interactions in NF54CSAh over NF54. Centromeric bins (gray) and *var* gene-containing bins (green) are annotated for each chromosome. (**N**) Differential interchromosomal interaction heatmap with intrachromosomal interactions removed.

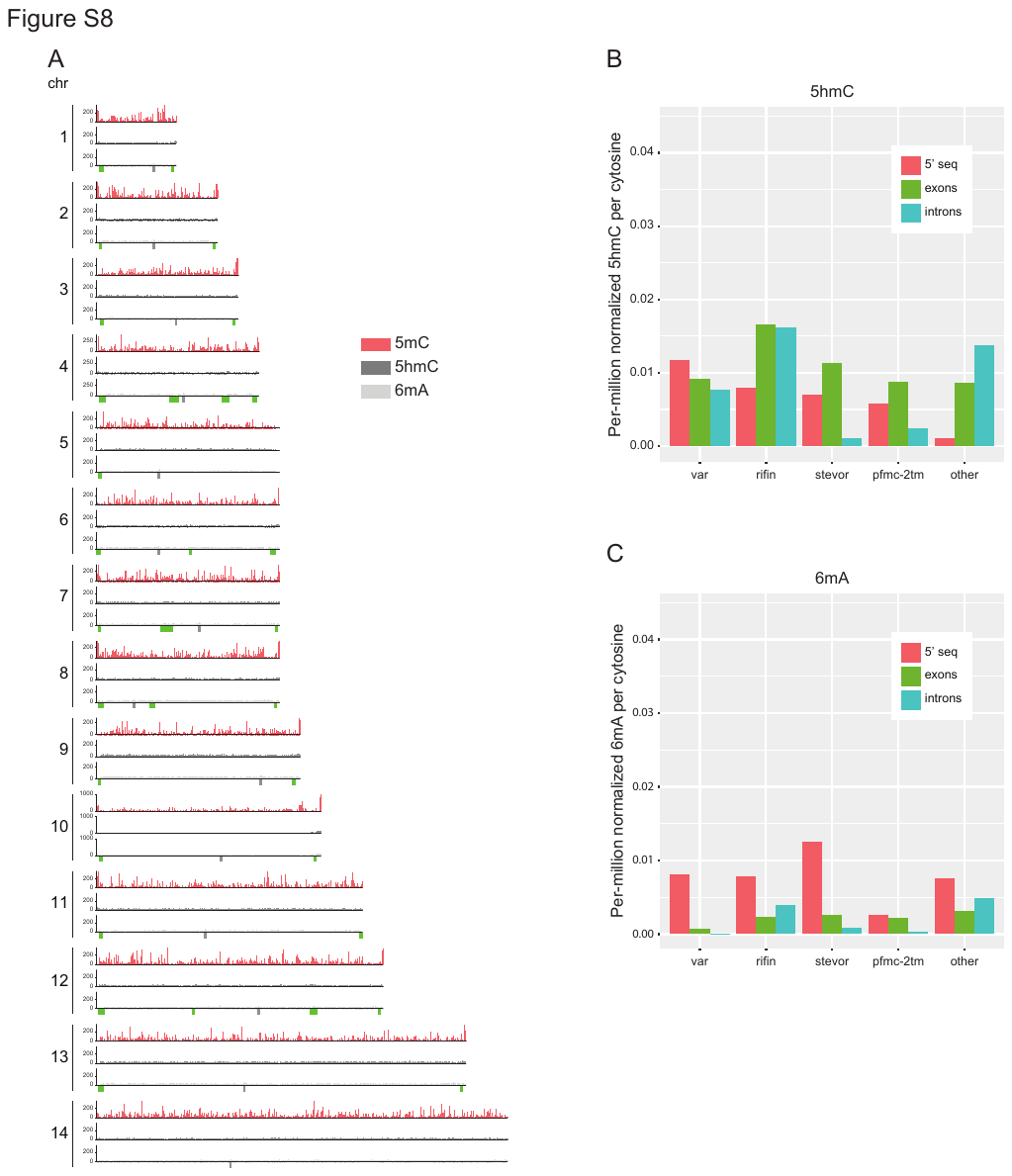

**Figure S8.** The P. falciparum genome contains very little 5hmC and 6mA compared to 5mC and no pattern between gene type or feature. (**A**) Genome-wide distribution of input-normalized 5-methylcytosine (5mC, top), 5-hydroxymethylcytosine (5hmC, middle) and 6-methyladenosine (6mA, bottom), with centromeres (gray) and *var* genes (green) annotated within each track. (**B and C**) Bar graphs depicting per-million read count normalized 5mC and 6mA per cytosine partitioned into the 500 bp 5’ region, exons and introns for four antigenically variable gene families (*var*, *rifin*, *stevor* and *pfmc-2tm*) as well as the remaining protein coding genes and intergenic regions.
